# The role of *GmXTH1*, a new xyloglucan endotransglycosylase/hydrolase from soybean, in regulating soybean root growth at seedling stage

**DOI:** 10.1101/2022.07.15.500193

**Authors:** Yang Song, Ye Zhang, Ye-yao Du, Sujie Fan, Di Qin, Zhuo Zhang, Pi-wu Wang

## Abstract

Soybean is an important crop for both food and oil, providing high quality protein and oil for humans and animals. Drought stress has an significant effect on the yield and quality of soybean. In the early vegetative stage of soybean, the developed root system can effectively improve the tolerance of soybean plant to drought. Xyloglucan endoglycosylase/hydrolase (XTH) is a cell wall remodeling enzyme involved in cell wall expansion and degradation. However, there are few studies on XTH’s regulation of root growth and response to drought stress in soybean seedlings.The objectives of this study were to clone GmXTH1 gene and determine the effects of over and interference-expression of GmXTH1 on soybean root development at seedling stage. A 1,015 bp full-length of the GmXTH1 gene was cloned from soybean (Glycine max L. Merril) cv. RM18 with a 840 bp open reading frame (ORF) of a putative protein. The over and interference expression vectors of GmXTH1 were transformed into soybean using Agrobacterium method. Transformed strains OEA1, OEA3, IEA4, IEA5 were identified by molecular detection methods. The roots of transformed strains OEA1and OEA3 with GmXTH1 gene overexpression are clearly developed and the root phenotypic attributes, including main root length, primary root number, lateral root length, root surface area, root volume, root dry weight, root fresh weight are higher than the control. In contrast, root attributes of IEA4 and IEA5 with GmXTH1 gene interference expression were significantly lower than those of the control, suggesting that the expression of GmXTH1 gene has an important effect on the soybean root growth and development at seedling stage. Under drought stress conditions, the activities of Superoxide dismutase (SOD), Peroxidase(POD)and Catalase(CAT)in the transgenic strains with an overexpression of the GmXTH1 gene were significantly increased, and the root activity, leaf relative water content and total chlorophyll content were ranked in order of OEAs, CK, and IEAs. In conclusion, GmXHT1 plays an important role in the growth of soybean root at seedling stage. Overexpression of GmXTH1 gene can promote root development of soybean seedling. The results of activities of protective enzymes (SOD, POD, CAT), root activity, leaf relative water content and total chlorophyll content under drought stress indicated that transgenic strains OEA1 and OEA3 with overexpressing GmXTH1 gene were resistant to drought stress, suggesting that GmXTH1 may play a positive role in responding to drought stress at soybean seedling stage.

## Background

Soybean [Glycine max L. (Merr.)] is one of the most important sources of oil and protein in the human diet and in animal feed. It is also the main economic and oil-bearing crop, and its yield is highly affected by soil water supply^[1]^. Root systems are the primary site of water absorption and play an important role in drought stress resistance and recovery^[2]^. Soybeans can survive drought stress if there is a robust and deep root system at the early vegetative growth stage. Numerous research publications have made it quite clear that a better understanding of the molecular and genetic basis for variation in root system will greatly aid the development of crop varieties with improved and more efficient nutrient and water acquisition under limiting conditions^[3]^. There are many studies on the cloning of soybean drought resistance genes, but the research on the cloning and drought resistance of soybean *XTHs* gene has not been reported so far.

Cell growth is an important aspect of plant growth and development, in the process, accompanied by cell wall remodeling^[4-5]^. Xyloglucan is the most important hemicellulose in the primary cell wall of dicotyledonous and non-grass monocots. Xyloclucan endo-transglucosylase/hydrolase (XTH) is a kind of cell wall relaxation enzymes, widely present in various tissues and cells of plants, modify the cellulose-xyloglucan complex of plant cell walls by catalyzing the cleavage and re-ligation of xyloglucan molecules, achieve cell wall remodeling. Studies have shown that it plays a vital role in plant cell wall remodeling process ^[6-8]^, in response to plant hormones and other environmental signals, closely related to plant growth and development, and regulated in plant growth on playing an important role in the physiological process.

XTH is a type of cell wall relaxation enzyme belonging to the subfamily of the glycoside hydrolase family GH16 family^[9]^. Higher plants usually contain 20 to 60 XTH proteins [10]. XTHs have 33 members in the Arabidopsis genome, 29 members in rice, 41 members in Populus euphratica and 25 members in tomato ^[10-14]^. According to the sequence characteristics of XTHs, they are divided into three categories: I, II and III, and class III XTHs are divided into two subclasses of IIIA and IIIB ^[15]^. In recent years, XTHs of class I, II and IIIB have been found to have significant xyloglucan endotransglucosylase (XET) activity, which is capable of endo-cutting xyloglucan molecules and rejoining the resulting reducing ends. In another xyloglucan chain, the class IIIA is characterized by xyloglucan endohydrolase (XEH) activity, specifically hydrolyzing xyloglucan β-1,4 glycosidic bonds. Moreover, members of different XTH subfamilies exert different activities. For example, AtXTH27, SlXTH5 and HvXTH8 belong to class IIIB and mainly exert XET activity ^[16-18]^. Many members of the I/II subfamily of XTHs proteins, such as AtXTH26, AtXTH14, AtXTH22, PttXET16-34, have only XET activity ^[19]^. The XTHs enzyme contains the characteristic motif DEIDFEFLG, which contains amino acid residues that mediate catalytic activity. N-glycosylation of serine or threonine residues near this catalytic site is also important for enzyme activity ^[20]^. The N-glycosylation site is conserved in the XTHs protein class I/II, but this site is not found in the IIIA subclass ^[21]^. The C-terminus of the XTHs protein often contains highly conserved cysteine, which can form The disulfide bond facilitates the stability of the XTH protein structure.

The xyloglucan endoglycosidase/hydrolase (XTH) catalyzes the cleavage and re-ligation of xyloglucan molecules. The mechanism of action of this enzyme is: XTH enzyme binds to the substrate which is hydrolyzed to form a glycosyl-enzyme covalent intermediate, which in turn transfers the glycosyl group to the reducing end of the polysaccharide residue for transglycosylation (XET activity); or transfers the glycosyl group to a water molecule for hydrolysis (XEH activity)^[22-24]^. The most suitable substrates for XTH are known as xyloglucan and xylooligosaccharides. The xylose side chain is an essential structure for the XTH receptor substrate, and XTHs with xyloglucan endosynthase activity are capable of accepting xyloglucan oligosaccharides. The three-dimensional structure of the protein shows that there is a 35Å gap in the XTH active site that can accommodate 7 main chain glucose groups. The -1 site unsubstituted glucosyl group plays an important role in maintaining the conformation of the enzyme-bottom complex as a catalytic reaction of important conditions ^[25-28]^. The xylose substitution at the ^+2^ subsite also has a very important effect on the enzyme activity, which may be an important recognition site for the enzyme ^[29]^; In addition, galactosyl or fucose side chains can also be to some extent Affect the activity of XET ^[30-32]^.

In recent years, *XTH* gene has been cloned from Arabidopsis thaliana, Gerbera jamesonii, Rosa chinensis, and other ornamentals^[33]^. But the related studies are still not enough in soybean. In this study, the new gene *GmXTH1* derived from soybean roots was cloned by genetic engineering technology. The function of target gene was identified on the effects of responding to drought stress and regulating the growth and development of soybean seedling roots by over-expressing and interference-expressing the *GmXTH1* gene to lay the theoretical basis for regulating the growth and development of soybean roots by molecular biology, and provide new genetic resources and basic materials for breeding soybean varieties with developed roots.

## Results

### Cloning of *GmXTH1* gene full-length cDNA

The conservative fragment of *GmXTH1* was cloned by reverse transcription PCR (RT-PCR) (Figures S1, A). The 3’RACE and 5’RACE procedures yielded nucleotide sequences of 422bp and 632bp, respectively (Figures S1, B, C). The sequencing results were aligned, and the duplicated sequences were removed. A specific primer to amplify the full-length gene product was designed, and the full-length cDNA, 1015 bp in length, was obtained via RT-PCR (Figure S1, D). To differentiate this gene from the known soybean *XTH23-like* gene, the target gene was named *GmXTH1*.

### Bioinformatics analysis

Based on online analyses, the complete open reading frame (ORF) of the target gene was determined to be 840bp, encoding 279 amino acid. The fragment of open reading frame on *GmXTH1* gene was obtained by PCR (Figure 1, A). The analysis of ProtParam showed that the molecular weight of the expressed protein was 31286.9 Da. The theoretical isoelectric point was 6.74. The total number of positively charged amino acid residues (Arg + Lys) is 24. The total number of negatively charged amino acid residues(Asp + Glu) is 25. Molecular formula is C_1374_H_2064_N_394_O_415_S_17_. Amino acid group analysis showed that the content of serine (Ser), glycine (Gly), threonine (Thr) were 9.7%, 8.6%, 7.5% respectively, which were higher than the others. The grand average of hydropathicity (GRAVY) is -0.458.

**Fig.1.**
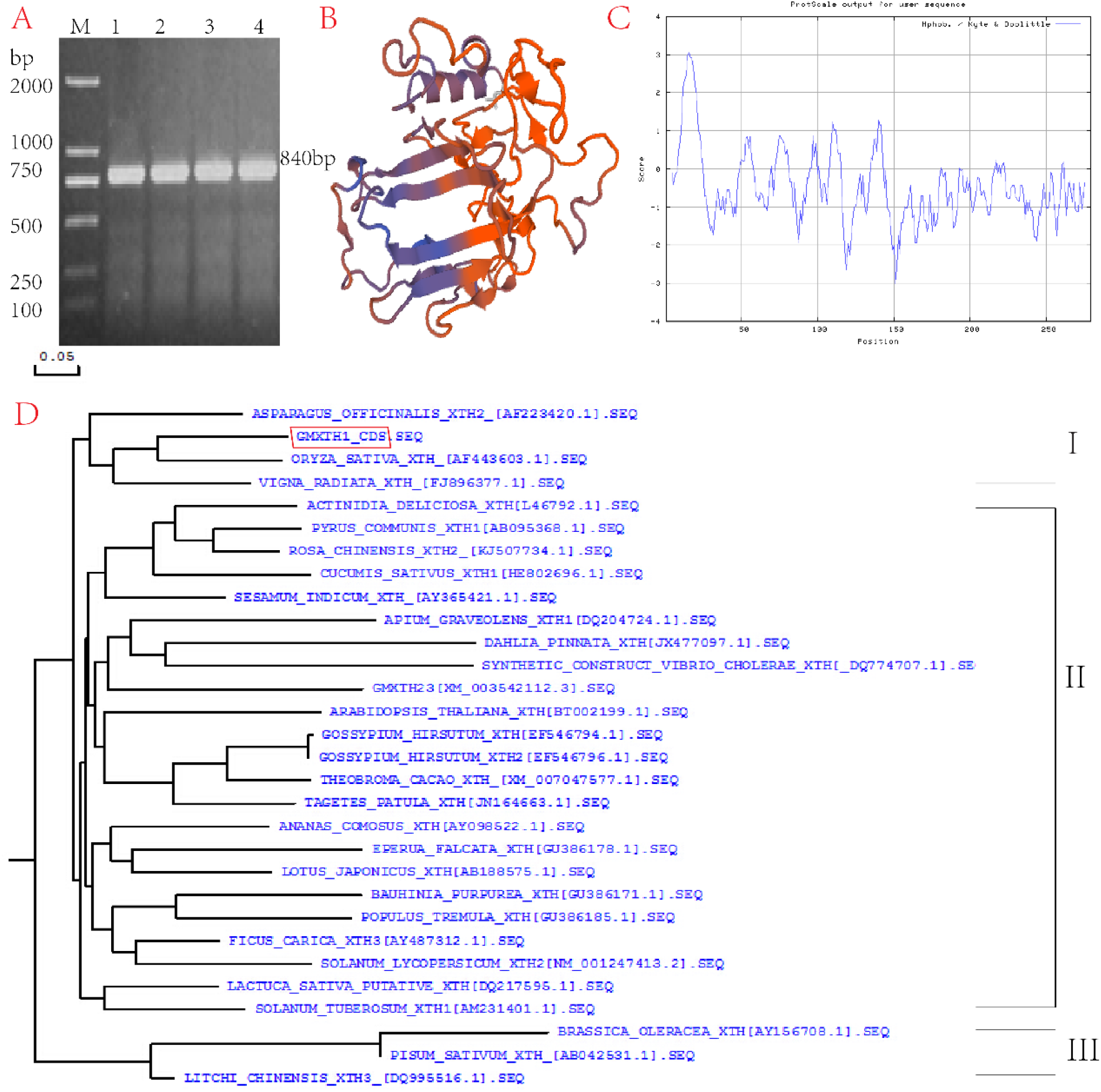
Bioinformatics analysis of *GmXTH1*. A. PCR product of open reading frame on *GmXTH1* gene; B. dimensional structure of GmXTH1 protein; C. hydrophobicity-hydrophilicity prediction of GmXTH1 protein; D. phylogenitic tree of GmXTH1 protein A. M: DNA Marker; 1-4: PCR product

The prediction results of Protein hydrophilic-to-hydrophobic showed that valine (Val) at 16-bit has the highest score (3.056) with strong hydrophobicity, and arginine (Arg) at 151-bit has the lowest score (−3.011) with strong hydrophilic. On the whole, there are more GmXTH1 hydrophilic amino acids than hydrophobic amino acids. Therefore, GmXTH1 is hydrophilic (Figure 1, B).

After analysis with DNAMAN software, the target gene *GmXTH1* and the known soybean gene *GmXTH23-like* were found to have 51.48 % homology at the nucleotide sequence level and 46 % at the amino acid sequence level. The amino acid sequences of all of the known XTHs in leguminous plants were downloaded from the NCBI website, and XTH protein phylogenetic trees were constructed using MEGA4.The neighbor-joining method was used to divide the Evolutionary analysis results into three groups(Groups I–III) based on the sequence homology of the XTHs (Figure 1, C). The soybean GmXTH1 showed high homology to Group I and a shorter genetic distance to both the Oryza Sativa XTH and Vigna Radiata XTH than to the soybean GmXTH23-like, indicating that the evolutionary history of GmXTH1 differs from that of GmXTH23-like.

### Construction of plant expression vectors

The plant over-expression vector pCAMBIA3301-GmXTH1 was successfully obtained using the restriction enzymes *Bgl*II and *BstE*II and confirmed by PCR as well as restriction enzyme digestion (Figure 2, A). The *GmXTH1* was amplified to obtain 840bp fragments by PCR, the pCAMBIA3301 vector and *GmXTH1* fragment of 840bp were obtained by double digestion, indicating that the *GmXTH1* was integrated into pCAMBIA3301 (Figure 2, C) and the plant over-expression vector pCAMBIA3301-GmXTH1 was constructed successfully.

**Fig. 2.**
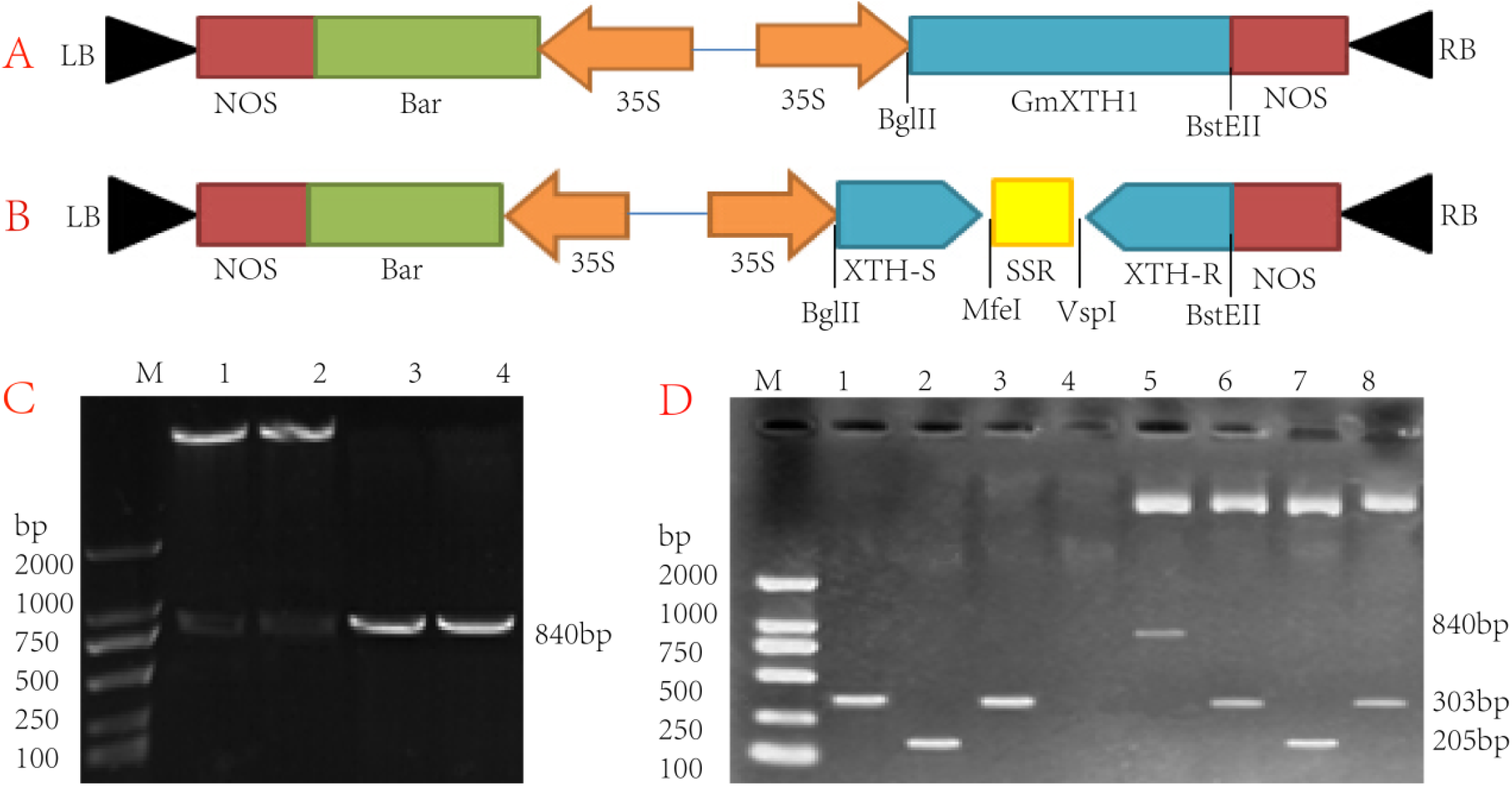
Construction of plant expression vectors. A. T-DNA structural region of plant over-expression vector; B. T-DNA structural region of RNA interference-expression vector; C. verification of plant over-expression vector; D. verification of plant RNA interference-expression vector C. M: DNA Marker; 1-2: double enzyme product by *Bgl*II and *BstE*II; 3-4: PCR product D. M: DNA Marker; 1: PCR product of *XTH* sense fragment; 2: PCR product of SSR fragment; 3: PCR product of *XTH* anti-sense fragment; 4: Blank channel; 5: Double enzyme product of *Bgl*II and *BstE*II; 6: Double enzyme product of *Bgl*II and *Mfe*I; 7: Double enzyme product of *Mfe*I and *Vsp*I; 8: Double enzyme product of *Vsp*I and *BstE*II

The plant RNAi-expression vector pCAMBIA3301-GmXTH1-RNAi was successfully obtained using the restriction enzymes *Bgl*II and *Mfe*I, *Mfe*I and *Vsp*I, *Vsp*I and *BstE*II, respectively. The XTH-sense, intron-SSR, XTH-antisense were connected into the vector by T_4_ DNA Ligase (Figure 2, B). The fragments of XTH-sense, intron-SSR, XTH-antisense were amplified to obtain 303bp, 205bp, 303bp fragments by PCR, respectively. The pCAMBIA3301 vector and XTH-sense, intron-SSR, XTH-antisense and XTH-RNAi fragments of 303bp, 205bp, 303bp, and 811bp were obtained by double digestion (Figure2, D), indicating that the plant RNAi-expression vector pCAMBIA3301-GmXTH1-RNAi was constructed successfully.

### Molecular detection of transgenic plants

A total of 19 resistant seedlings were obtained using an Agrobacterium-mediated method, of which five transgenic over-expression vector plants were PCR positive for the target fragments of *GmXTH1* and *Bar*(Figure 3, A), and six transgenic RNAi-expression vector plants were PCR positive for the target fragments of *Bar* and *35S*(Figure 3, B). Matured seed were harvested from positive T0 plants and planted in a greenhouse at room temperature. Transgenic plants from the each event at T1 and T2 generations were detected by PCR for the *GmXTH1, Bar*, and *35S* target fragments. Five positive transgenic strains (named OEA1-OEA5) with plant over-expression vector and five positive transgenic strains (named IEA1, IEA3-IEA6) with plant RNAi-expression vector were obtained.

**Fig.3.**
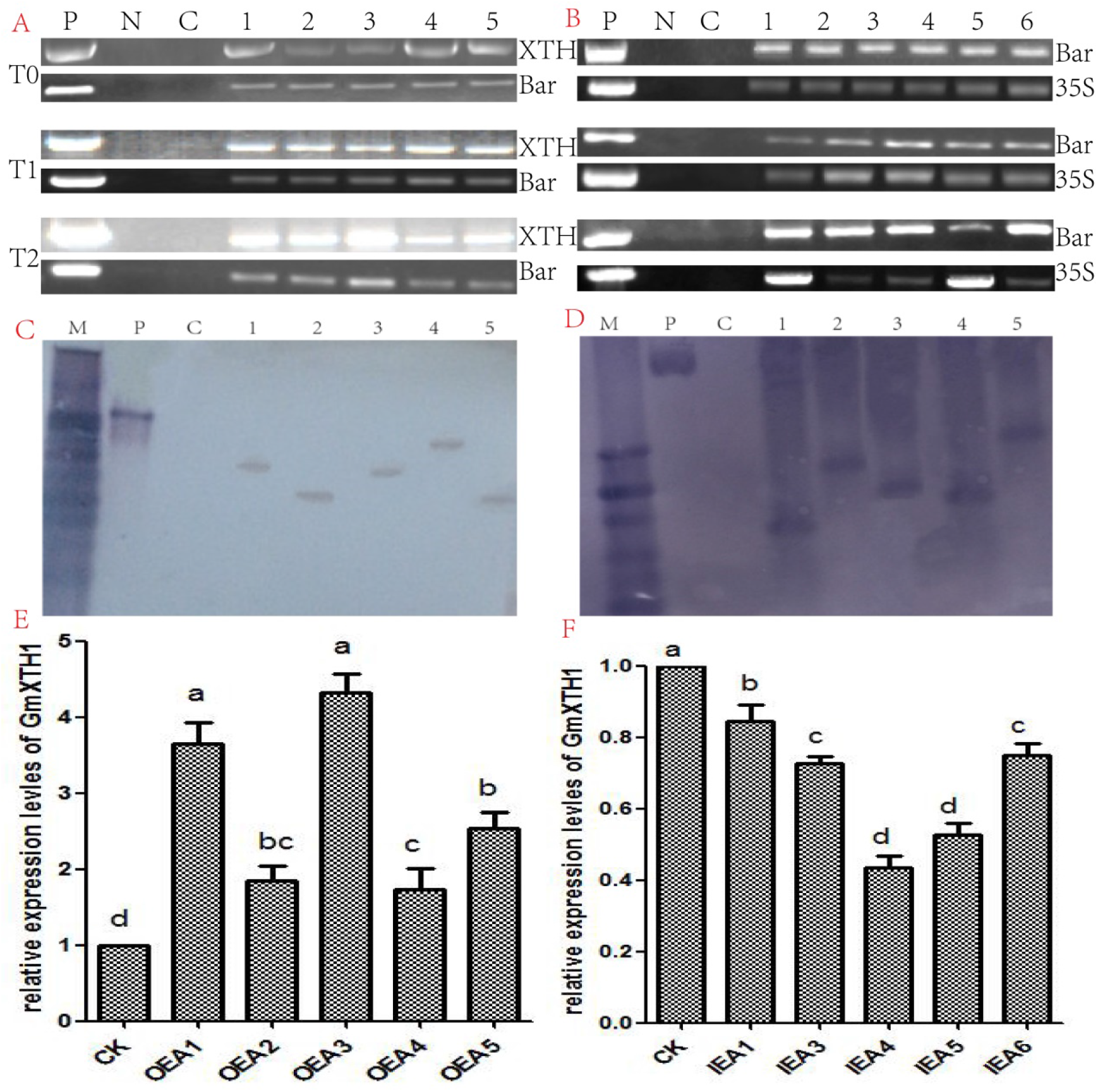
Molecular detections of transgenic plants. A. PCR detections of transgenic plants with plant over-expression vector at T0-T2 generations; B. PCR detections of transgenic plants with plant RNA interference-expression vector at T0-T2 generations; C. Southern blot analysis of transgenic plants OEAs; D. Southern blot analysis of transgenic plants IEAs; E. relative expression levels of GmXTH1 in transgenic plants OEAs; F. relative expression levels of GmXTH1 in transgenic plants IEAs M :DNA Marker;P :Positive control of plasmid ;N :Negative control of water;CK : Negative control of transformed plant;1-5 :Putative transgenic plants

To further confirm the presence of transgenes in the T2 generation, a southern hybridization procedure was performed on five transgenic plants from the T2 generation that had positive PCR results. Strong hybridization signals were observed for *Bar* selection marker gene to OEA1-OEA5 and IEA1, IEA3-IEA6, respectively, indicating that the foreign genes were successfully integrated into the soybean genome. The sizes of the hybridized fragments indicated that the insertion sites of the target genes were different, and the single fragment implied that one copy was inserted (Figure 3, C, D).

According to the quantitative real time-PCR results (Figure 3, E, F), the transcript levels of *GmXTH1* showed large differences among the different transgenic strains. The experimental results were analyzed with the 2^-ΔΔCt^ method, and the expression of *GmXTH1* in the root of control plants was set to 1. Compared with the control plants, the relative expression level of *GmXTH1* gene was significantly increased in transgenic lines with over-expression of *GmXTH1* gene. The average expression of *GmXTH1* gene was the highest in the OEA3, which was 4.3327 times of that of the control, followed by the OEA1, which was 3.66403 times than the control (Figure 3, E). In contrast, the relative expression level of *GmXTH1* gene was significantly decreased in the transgenic lines with interference-expression of *GmXTH1* gene, and the decrease of IEA4 was the highest, which was 56.3163%, followed by IEA5, which was 47.303% lower than that of control (Figure 3, F).

### Effects on the root development of transgenic plants with different expression of *GmXTH1* gene

The apparent morphological observation of transgenic strains’ roots showed that the roots of transgenic plants with *GmXTH1* gene over-expression are more developed than the control and the ones with *GmXTH1* gene interference-expression (Figure S2). The results of scanning detection on roots showed that the mean values of main root length, number of primary lateral root, lateral root length, root surface area, root volume, root dry weight and root fresh weight of over-expression lines OEA1 and OEA3 were significantly higher than the control, indicating that the over expression of *GmXTH1* gene significantly promoted the growth and development of soybean roots at seedling stage, and the mean value of root phenotype was increased significantly. However, the average of these indicators of interference -expression lines IEA4 and IEA5 is obviously lower than the control, suggesting that the expression level of *GmXTH1* gene has an important effect on the growth and development of soybean roots at seedling stage (Table 1).

**Table.1.**
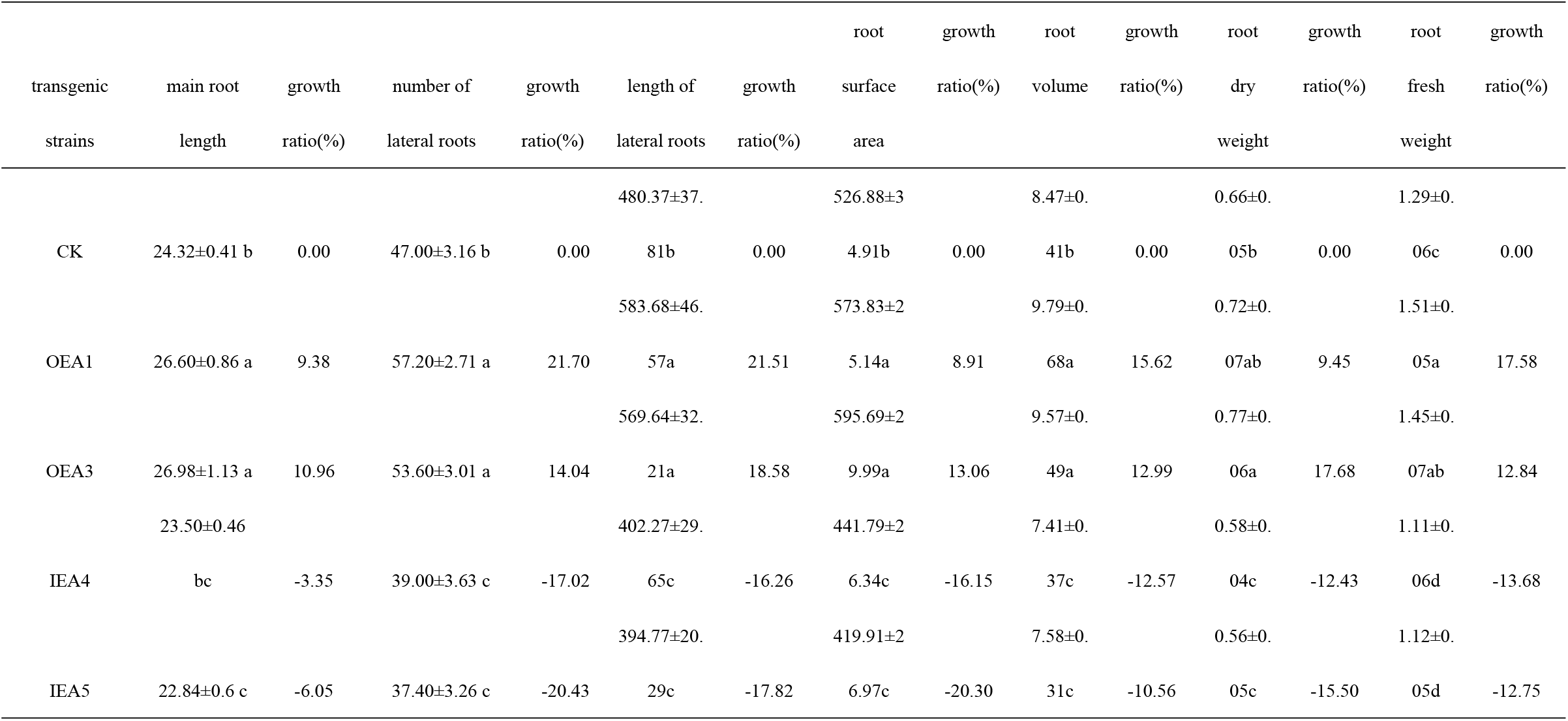
Root phenotypic traits of different transgenic plants

### Effects of drought stress on antioxidant enzyme activities of transgenic plants

According to the SOD activity measurement results, under normal condition, the SOD activity of over-expression lines OEA1 and OEA3 was significantly higher than that of the control. On the contrary, the SOD activity of IEA4 and IEA5 was significantly lower than that of control. Under the drought stress condition, the SOD activity of the control, OEAs and IEAs was significantly increased (P <0.01). The mean activity of SOD activity was as follows: OEAs> CK> IEAs. The SOD activity of OEA3 with GmXTH1 gene over-expressing was the highest, indicating that OEA3 had stronger ability to resist reactive oxygen species and free radical damage (Figure 4, A). As shown in Figure 4B, the difference of leaf POD activity between OEAs, IEAs and CK was not significant under normal condition. Under drought stress condition, the POD activity of OEAs, IEAs and CK was significantly increased, in which OEA1 and OEA3 were respectively 18.39% and 17.08% higher than the control, IEA4 and IEA5 were 2.8% and 1.19% lower than control.

**Fig.4.**
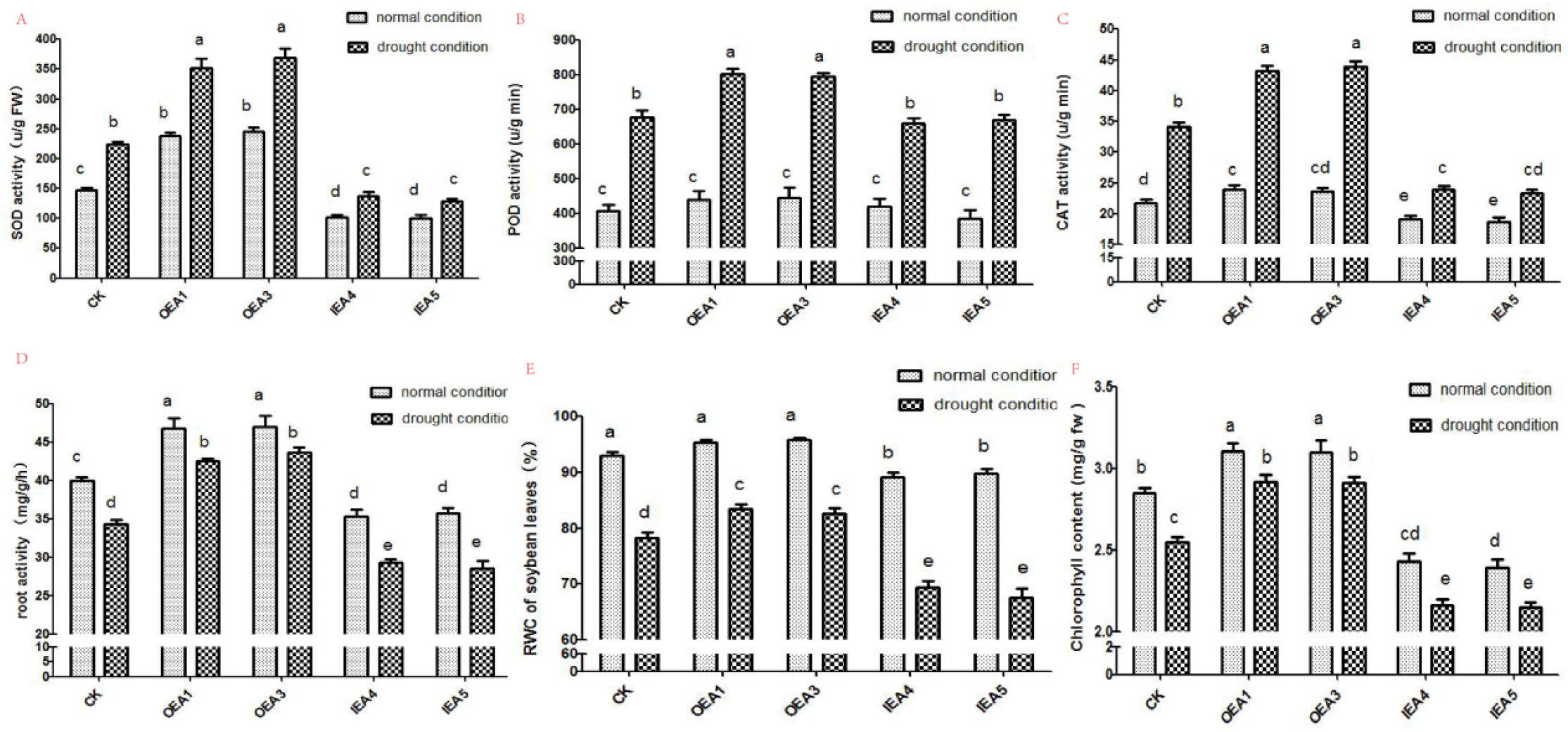
Changes in physiological and biochemical indexes of different transgenic strains under drought stress.

The mean activity of CAT activity was as follows: OEAs> CK> IEAs under normal condition. The CAT activity of OEAs, IEAs and CK was significantly increased under drought stress condition. The variation trend of different strains is quite different, which the average activity of OEA1 and OEA3 strains was 26.19% and 28.36% higher than that of the control, respectively. Conversely, the average CAT activity of IEA4 and IEA5 strains decreased by 29.93% and 31.77% compared with the control (Figure 4, C).

Under normal water treatment, the average of root activity was expressed as OEA3> OEA1> CK> IE5> IE4, and the difference was significant between OEAs, IEAs and CK, indicating that the expression of *GmXTH1* gene significantly affected the root activity of soybean seedling. Under the condition of drought treatment, root activity of soybean plants showed a downward trend, and the average root activity was OEA3> OEA1> CK> IE4> IE5. The average root activity was as follows: OEAs> CK> IEAs whether under normal water condition or drought stress condition, indicating that the transgenic strains with *GmXTH1* gene over-expression had a higher root activity with stronger drought resistance(Figure 4, D).

As shown in Figure 4E, the leave relative water content (RWC) was expressed as OEAs> CK> IEAs, of which IEAs was significantly lower than that of the control and OEAs. Under drought stress, the RWC of each strain was significantly decreased, which expressed as OEAs> CK> IEAs with a significant difference. This result showed that the OEAs’ RWC was relatively much higher than the others no matter under normal condition or drought condition, indicating that transgenic strains with over expression of *GmXTH1* gene may increase the water retention of soybean seedlings compared with CK and IEAs.

Chlorophyll plays a crucial role in photosynthesis as an important pigment. According to the Figure 4F, the total chlorophyll content was expressed as OEAs> CK> IEAs whether it is drought stress or normal water treatment. The leaf chlorophyll content of soybean seedling under drought stress was significantly lower than that of normal treatment, and the decrease of OEAs (18.708-18.687%) was less than that of IEAs (24.512-26.728%).

### Investigation of agronomic traits in field on transgenic strains

Under the condition of normal field management, there was no significant difference between the transgenic strains and the control strains RM18 in leaf shape, flower color, coat color, growth period and pod habit, which were expressed as round leaves, purple flowers, brown hairs and infinite pod habit. There was no significant difference in plant height, pitch number, pod number, hundred-grain weight and seeds numbers per plant between transgenic strains and non-transgenic ones. Yield results are shown as OEA3> OEA1> CK> IEA5> IEA4, in which strain OEA3 increased significantly, the yield increased by 8.19%. The difference between the remaining materials did not reach a significant level.

## Discussion

Drought is one of the most common abiotic stresses limiting plant growth and crop production in many areas of China and in the rest of the world^[34-36]^.Soybean drought resistance is a comprehensive trait that the mechanism of drought resistance is complex^[37]^. Therefore, the evaluation of plant drought resistance requires multiple indicators of comprehensive analysis of the results obtained more reliable. In this study, the transgenic soybean lines overexpressing GmXTH1 gene and interfering with GmXTH1 gene were studied in the seedling stage, and the SOD, POD, CAT activity, root activity, leaf water loss rate and chlorophyll content of soybean seedlings were observed. The changes of these indicators under drought stress were used to evaluate the drought resistance of different strains at the seedling stage.

Plant roots are active absorbing organs and synthetic organs. The growth and vitality level of roots directly affect the nutritional status and yield level of aboveground roots^[38-39]^. Root activity is a quantity that characterizes plant roots. Its size can judge the adaptability of plants and is relatively stable. It can be used as the main indicator in the test^[40]^. The results of this study indicated that the expression of GmXTH1 gene significantly affected the root activity of soybean seedlings. The overexpressing lines had higher root activity and stronger drought resistance. On the contrary, the interference expression of GmXTH1 gene inhibited the development of soybean roots at the seedling stage, and the indexes were significantly lower than the control, suggesting that the expression of XTH gene has an important influence on the growth and development of soybean roots at seedling stage.

The relative water content (RWC) of plant leaves is an important indicator to reflect the water physiological state of plant leaves. When the plants are stressed by stress, the size of RWC in leaves can reflect the water deficit of plant tissues to a certain extent, and it can also reflect its resistance^[41-42]^. The inverse strength, relatively high RWC can effectively maintain the structure of the chloroplast and protect the photosystem II (PSII), and improve the effective photosynthesis of plants^[43]^. The results of this study showed that under drought stress, the leaf water content of each line decreased significantly. Under normal conditions and under drought conditions, the leaf water content of the overexpressed lines was relatively high, indicating overexpression of GmXTH1. The gene can maintain the leaf water content of soybean seedlings at a high level, which is beneficial to maintain the integrity of its chloroplast structure and PSII function, so that it maintains a high level of photosynthesis. It can be seen that the drought resistance of overexpressing lines is stronger than that of the control. M18 and interference expression lines.

Chlorophyll is an important component of chloroplasts, which directly affects the absorption, transmission and transformation of photons in photosynthesis. It is the material basis for plants to absorb solar energy for photosynthesis. The results showed that the chlorophyll content of soybean seedlings under drought stress was significantly lower than that of normal treatment, indicating that drought treatment can inhibit chlorophyll synthesis and accelerate its decomposition, which may be related to drought stress, which causes chloroplast to produce membrane lipid peroxidation. Malondialdehyde is associated with the destruction of chlorophyll. However, whether it was drought treatment or normal water treatment, the chlorophyll content showed GmXTH1 gene overexpression strain>control M18>GmXTH1 gene interference expression strain, indicating that the leaves of overexpressing lines had higher light energy absorption in photosynthesis. And transformation, indicating that the overexpressing strain has stronger drought resistance than the control M18 and the interfering expression strain.

In the soybean seedling stage, the GmXTH1 gene was overexpressed, the GmXTH1 gene was interfered with expression, and the control mutant material M18 was treated with normal water treatment and drought stress, and comprehensive protective enzyme activity, root activity, relative water content of leaves, chlorophyll, etc. The index concluded that the transgenic lines overexpressing the GmXTH1 gene had significantly higher drought resistance than the control M18. Conversely, the GmXTH1 gene-interfering transgenic lines were less resistant to drought than the control. Moreover, the growth status of roots of overexpressing lines at seedling stage was significantly better than that of control M18, and the agronomic traits did not change significantly compared with the control. Transgenic lines overexpressing GmXTH1 gene, OEA1 and OEA3, have good agronomic traits and strong drought resistance, and can be used as a basic material for soybean drought-resistant breeding.

## Materials and Methods

### Plant materials, growth conditions, and drought stress treatment

Seeds of soybean RM18 (a stable root mutant strain with well developed root system) were provided by the Center for Plant Biotechnology, Jilin Agricultural University. Plants were grown in the cylindrical plastic tube with diameter 7cm and height 22cm using sanding method at greenhouse (16 h light, 25°C/8 h dark, 22°C photoperiod). The plants were subjected to water stress by repeated drought method. After 10 days of drought treatment, the water was rehydrated, making soil water content maintained at 80% ± 5%.

### Isolation and cloning of *GmXTH1*

The RNA of soybean seedling roots were extracted using an RNAiso kit (Takara, Otsu, Japan) according to the manufacturer’s instructions. The transcript levels of relevant genes were determined by real-time PCR. RNase-free DNase I was used to digest and treat the RNA to prevent DNA contamination.

After RNA extraction, the corresponding cDNA was synthesized using a RevertAid First Strand cDNA Synthesis Kit (Fermentas Company, Hanover, MD, USA). Based on the sequence of preliminary RNA-seq results, the nested primers XTHgsp were designed to amplify the target fragment by RT-PCR (Table S1). The target sequence was then cloned and sequenced to verify that the correct target fragment had been amplified.

The 3’end of the cloned fragment was amplified with a 3’RACE kit (Takara). RNA from fresh soybean roots was used as the template, and the 3’RACE adaptor primer from the kit was used for the reverse transcription reaction to synthesize first-strand cDNA. The XTHgsps primer and 3’RACE outer primer were used for the outer PCR, and the XTHgspas primer and the 3’RACE inner primer were used for the inner PCR. Amplification was performed to obtain the 3’end fragment of the gene.

The 5’end of the cloned fragment was amplified with a 5’RACE kit (Takara). The 5’RACE adaptor in the kit was used to evaluate the mRNA, and the multistep enzyme was used to speed up the reaction. Random 9-mers were used for reverse transcription to synthesize cDNA. The XTHgsps primer and 5’RACE outer primer were used for the outer PCR. The reaction product was used as the template for the inner PCR with primer XTHgspas and the 5’RACE inner primer. Amplification was performed to obtain the 5’end fragment of the gene.

The 3’RACE and 5’RACE products were aligned, and specific primers were designed to amplify the full-length gene segment (Table S1). The cDNA from the total RNA from soybean root was used as the template, and the segment corresponding to the GmXTH1 gene was amplified by RT-PCR.

### Bioinformatics analysis of *GmXTH1*

The online analytical procedure ORF FINDER (http://www.ncbi.nlm.nih.gov/gorf.html) was used to perform readable frame analysis for the target gene sequence. DNAMAN software (Version 5.0, Lynnon Biosoft Company, Quebec, Canada) was used to translate the ORF of the target gene into the amino acid sequence and to perform multiple sequence alignment. ProtParam software (Http://www.Web.Expasy.Org/protparam/) was used to predict the protein sequence theory parameters. The protein three-dimensional structure of GmXTH1 gene was constructed by SWISS-MODEL online software. Protscale software (http://web.Expasy.org//Protscale/) was used to predict the hydrophobicity of the protein. The sequences of all of the known XTHs were downloaded from the NCBI website, and MEGA (Ver4.0; Tamura et al. 2007) software was used to analyze protein homology and create a phylogenetic tree.

### Construction of plant expression vectors and soybean’s transformation

The plant expression vector pCAMBIA3301 was digested with restriction endonuclease *Bgl*II and *BstE*II, and the original GUS gene of pCAMBIA3301 vector was replaced by the cloned *GmXTH1* full length gene using T4 ligase. The plant over-expression vector pCAMBIA3301-GmXTH1 was constructed with *CaMV35S* as promoter and *Bar* as selection maker gene.

In the same way, the original *GUS* gene of pCAMBIA3301 vector was replaced by GmXTH1 sense fragment + SSR intron + GmXTH1 antisense fragment, constructing the plant RNA interference expression vector pCAMBIA3301-GmXTH1-RNAi.

Then the plant over-expression vector pCAMBIA3301-GmXTH1 and plant RNA interference expression vector pCAMBIA3301-GmXTH1-RNAi were transformed into soybean RM18 using Agrobacterium-mediated method.

### Molecular detection of transgenic plants

PCR detection of transgenic events was sequentially performed on T0, T1 and T2 transgenic plants. Fresh leaves (0.1 g) were collected from each of the transformed plants at 25 days after planting and DNA was extracted from the leaves using plant genome extraction kit (CWBIO company, Beijing, China). Three pairs of oligonucleotide primers named XTHover, Bar and 35S were designed to target these three genes using the Primer 5.0 software (PREMIER Biosoft, Palo Alto, CA, USA) (Table S1).

Southern blot analysis was performed to ensure that the foreign gene *Bar* was present in randomly excised segments of the T2 transformed soybean genome by following the protocol from the DIG High Prime DNA Labeling and Detection Starter Kit I (Roche, Indianapolis, IN, USA). DNA was extracted from the plants with positive PCR results, digested with *Hind*III, and then transferred to a nitrocellulose membrane. The DIG High Prime DNA Labeling and Detection Starter Kit I was used for probe labeling, hybridization, and chromomeric processing according to the manufacturer’s instructions.

Total RNA was extracted from soybean roots of T2 positive transgenic plants using a total RNA extraction kit (Fermentas). The corresponding cDNA was synthesized according to the Revert Aid First Strand cDNA Synthesis Kit (Fermentas) operation instructions. Using the cDNA as a template, β-actin (Genebank no: NM 001252731.2) as a reference gene, the target fragment of *GmXTH1* was amplified according to the SYBR Premix Ex TaqTM Kit (Takara, Otsu, Japan) operation instructions. The relative expression levels of the target gene were analyzed by the 2^-ΔΔCt^ method, using the formula reported by Lee and Schmittgen.

### The phenotypic characterization of soybean roots at seedling stage

The soybean roots at the V3 stage of seedling were scanned and observed by EPSON scanner (Seiko Epson Corp, Tokyo, Japan). The images of soybean roots were saved after scanning and analyzed by WinRHIZO analysis program (version 4.0b, Regent Instruments Inc., Quebec, Canada 2000). The main root length, number of primary lateral root, lateral root length, root surface area, root volume, root dry weight and root fresh weight were measured and analyzed. The correlation analysis and variance analysis were performed by DPS data analysis software. The LSR multiple comparison and the difference significance test (α = 0.05) were performed on the mean of the parameters. Five plants of each transgenic positive strain were selected to be tested and 3 replicates for per plant.

### The determination of physiological and biochemical indexes

The activity of superoxide dismutase (SOD) was determined by nitrogen blue tetrazolium (NBT) spectrophotometric method. The activity of peroxidase (POD) was measured by guaiacol method. The activity of catalase (CAT) was determined by UV absorption method. The relative water content (RWC) was determined by weighing method. The chlorophyll content was determined by acetone colorimetry. The experimental data were analyzed by Excel2003 and DPS software, and the difference between them was significant. Graphical analysis and mapping were performed with GraphPad Prism 5 software.

## Supplementary Information

## Acknowledgement

Funding for this research was provided by the Natural Science Foundation of China (No. 31571689).

## Author contribution statement

Wrote first draft YS, ZY and PW; designed experimental work YZ, YS, YD, DQ,JQ and PW; investigation YZ; provided experimental materials YS, PW, JQ; analyzed data YZ; wrote original manuscript YZ, YS; wrote and edit review YS,YZ; visualization YZ; supervised the whole work YS; project administration JQ. All authors have read and agreed to the published version of the manuscript.

## Data availability

The materials presented in this article will be freely available to any researcher wishing to use them for non-commercial purposes, and the author responsible for distribution of materials in accordance with the policy described in the Instructions for Authors (www.springer.com) is: Pi-wu Wang (peiwuw@163. com).

## Declarations

### Ethical approval

The experiments reported in this study comply with the current laws of China.

### Open Access

This article is licensed under a Creative Commons Attribution 4.0 International License, which permits use, sharing, adaptation, distribution and reproduction in any medium or format, as long as you give appropriate credit to the original author(s) and the source, provide a link to the Creative Commons licence, and indicate if changes were made. The images or other third party material in this article are included in the article’s Creative Commons licence, unless indicated otherwise in a credit line to the material. If material is not included in the article’s Creative Commons licence and your intended use is not permitted by statutory regulation or exceeds the permitted use, you will need to obtain permission directly from the copyright holder.

## Supplement Tables and Figures

**Tab.S1.**
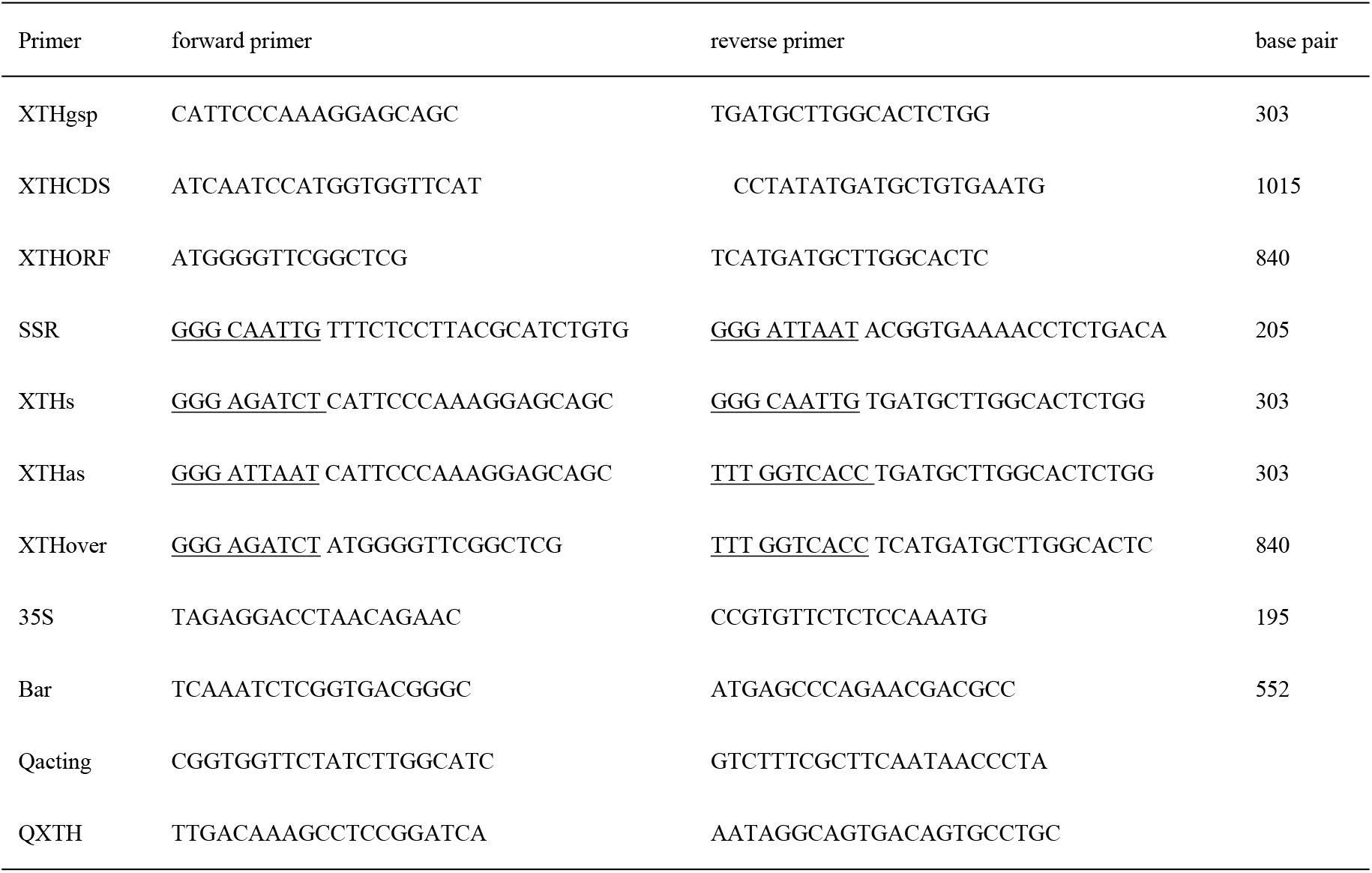
Primers for PCR

**Tab.S2.**
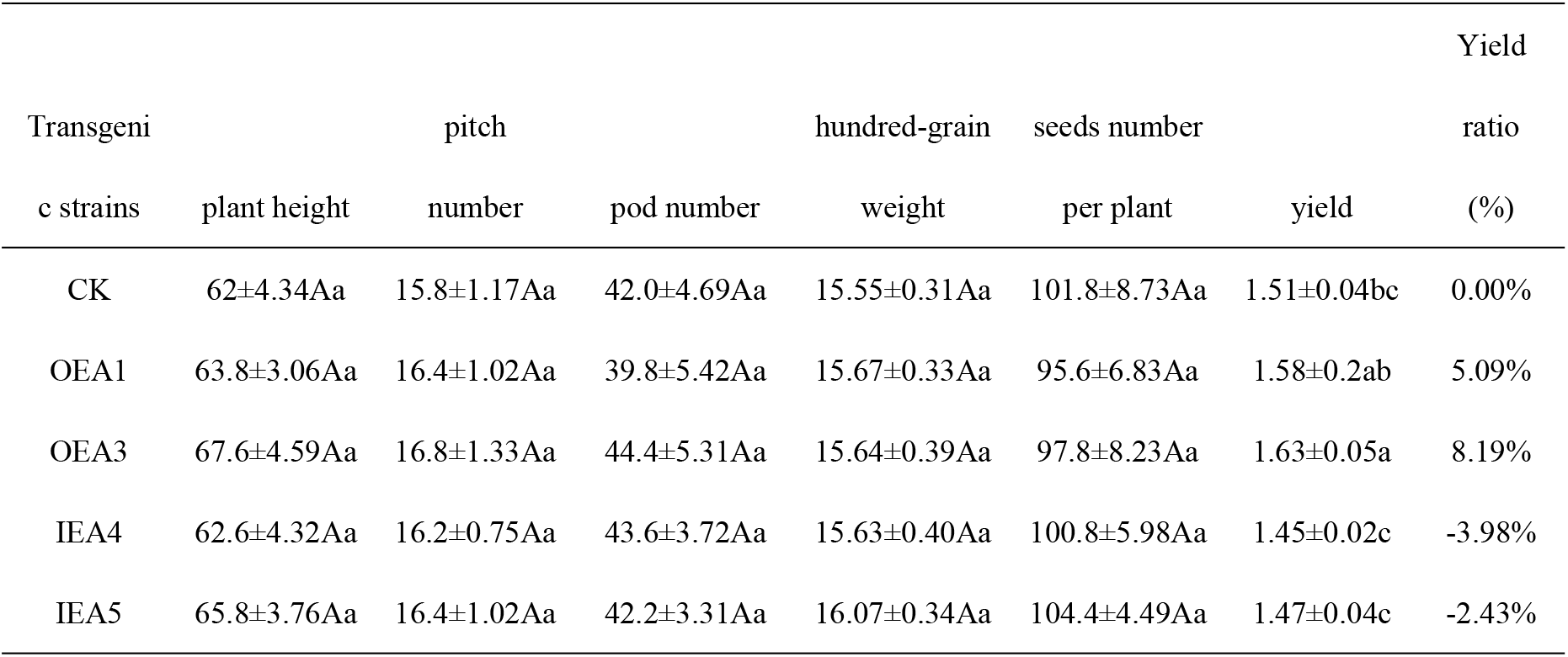
Yield traits investigation of transgenic soybean

**Fig.S1.**
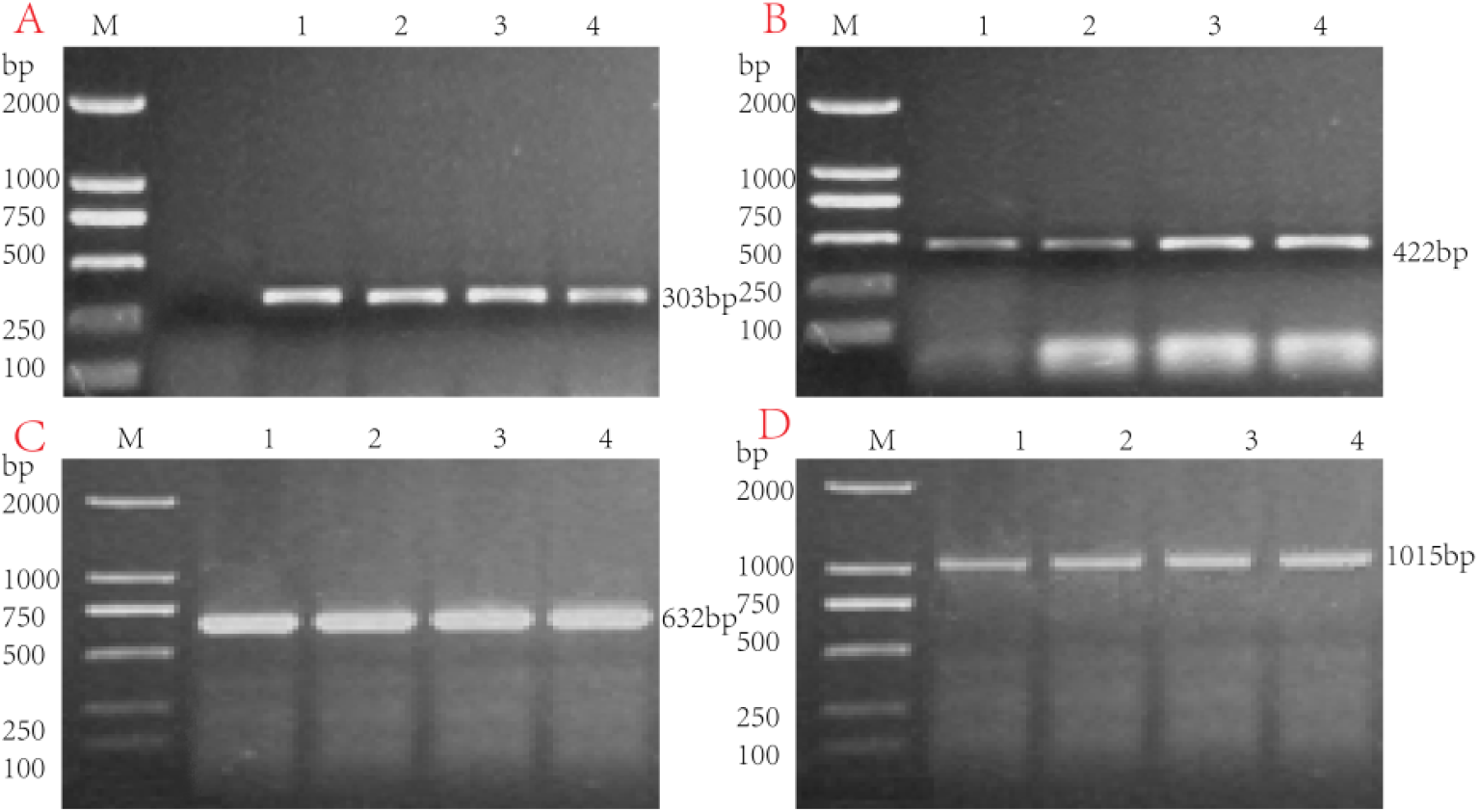
cloning of full-length *GmXTH1* cDNA. A. PCR product of conservative fragment on *GmXTH1* gene; B. PCR product of 3’
sfragment on *GmXTH1* gene; C. PCR product of 5′fragment on *GmXTH1* gene; D. PCR product of full *GmXTH1* gene M: DNA Marker; 1-4: PCR product

**Fig.S2.**
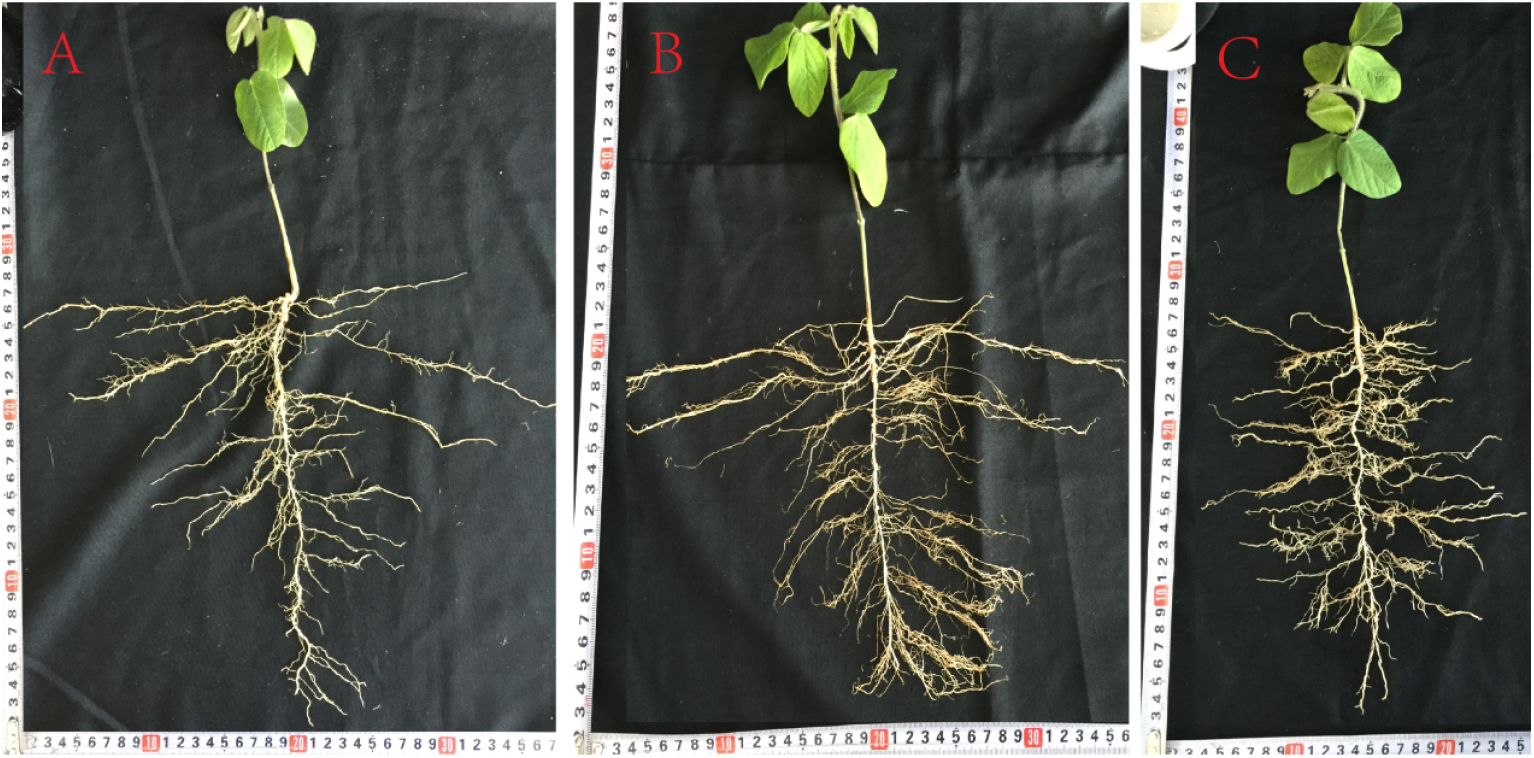
root performance of transgenic plants under normal condition. A. receptor; B. OE plant (transgenic plant with *GmXTH1* gene over-expression); C. IE plant (transgenic plant with *GmXTH1* gene interference-expression)

## References

[1] Diane Luth, Katey Warnberg, and Kan Wang. Soybean [Glycine max (L.) Merr.].2015,275–284.DOI:10.1007/978-1-4939-1695-5_22Corpus ID: 46381582

[2] Alok Ranjan,Ragini Sinha,Sneh L. Singla-Pareek,Ashwani Pareek,Anil Kumar Singh,Shaping the root system architecture in plants for adaptation to drought stress. Physiologia plantarum, 2022, 174(2):e13651,https://doi.org/10.1111/ppl.13651

[3] Md Nurealam Siddiqui, Jens Léon, Ali A Naz, Agim Ballvora, Genetics and genomics of root system variation in adaptation to drought stress in cereal crops, J Exp Bot. 2021, 24;72(4):1007–1019. doi: 10.1093/jxb/eraa487.

[4] Cosgrove D J. Growth of the plant cell wall [J]. Nat Rev Mol Cell Bio, 2005,6:850–861.

[5] Baumann M J, Eklof J M, Michel G, et al. Structural evidence for the evolution of xyloglucanase activity from xyloglucan endo-transglycosylases: biological implications for cell wall metabolism[J]. Plant Cell,2007,19(6):1947–1963.

[6] Saladié M, Rose J K C, Cosgrove D J, et al. Characterization of a new xyloglucan endotransglucosylase /hydrolase (XTH) from ripening tomato fruit and implications for the diverse modes of enzymic action[J].Plant J, 2006, 47(2):282–295.

[7] Sasidharan R, Chinnappa C C, Staal M, et al. Light quality-mediated petiole elongation in Arabidopsis during shade avoidance involves cell wall modification by xyloglucan endotransglucosylase/hydrolases[J]. Plant Physiol, 2010, 154(2):978–990.

[8] Soga K, Wakabayashi K, Kamisaka S, et al. Effects of hypergravity on expression of XTH genes in azuki bean epicotyls[J].Physiol Plantarum, 2007, 131(2):332–340

[9] Rose J K C, Braam J, Fry S C, et al. The XTH family of enzymes involved in xyloglucan endotransglucosylation and endohydrolysis: current perspectives and a new unifying nomenclature[J].Plant Cell Physiol, 2002, 43(12):1421–1435.

[10] Yokoyama R, Rose J K C, Nishitani K. A surprising diversity and abundance of xyloglucan endotransglucosylase/hydrolases in rice. Classification and expression analysis [J]. Plant Physiol. 2004,134:1088–1099.

[11] Geisler L J, Geisler M, Coutinho P M, et al. Poplar carbohydrateactive enzymes. Gene identification and expression analyses [J]. Plant Physiol, 2006, 140:946–962.

[12] Saladié M, Rose J K C, Cosgrove D J, et al. Characterization of a new xyloglucan endotransglucosylase /hydrolase (XTH) from ripening tomato fruit and implications for the diverse modes of enzymic action[J]. Plant J, 2006,47:282–295.

[13] Olsen S, Popper ZA, Krause K. Two sides of the same coin: xyloglucan endotransglucosylases/hydrolases in host infection by the parasitic plant cuscuta. Plant Signal Behav, 2016, 11: e1145336

[14] Albert M, Werner M, Proksch P, et al. The cell wall-modifying xyloglucan endotransglycosylase/hydrolaselexth1 is expressed during the defence reaction of tomato against the plant parasite cuscuta reflexa. Plant Biol, 2004, 6: 402–407

[15] Kallas A M,Piens K,Denman S E, et al. Enzymatic properties of native and deglycosylated hybrid aspen(Populus tremula× tremuloides)xyloglucan endotransglycosylase 16A expressed in Pichia pastoris[J].Biochem J, 2005, 390(1): 105–113

[16] Van Sandt V S T, Guisez Y, Verbelen J P, et al. Analysis of a xyloglucan endotransglycosylase/hydrolase(XTH)from the lyco-podiophyte Selaginella kraussiana suggests that XTH sequence characteristics and function are highly conserved during the evolution of vascular plants[J]. J Exp Bot,2006, 57(12): 2909–2922.

[17] Han Y, Wang W, Sun J, et al. Populus euphratica xth overexpression enhances salinity tolerance by the development of leaf succulence in transgenic tobacco plants[J]. J Exp Bot, 2013, 64: 4225–4238

[18] Matsui A, Yokoyama R, Seki M, et al. AtXTH27 plays an essential role in cell wall modification during the development of tracheary elements[J]. Plant Journal, 2005, 42(4):525–534.

[19] Liu Y B,Lu S M,Zhang J F, et al. A xyloglucanendotransglucosylase/hydrolase involves in growth of primary rootand alters the deposition of cellulose in Arabidopsis[J]. Planta,2007,226(6): 1547–1560.

[20] Cho S K,Kim J E,Park J A, et al. Constitutive expression of abioticstress-inducible hot pepper CaXTH3,which encodes a xyloglucan endotransglucosylase/hydrolase homolog,improves drought and salt tolerance in transgenic Arabidopsis plants [J]. FEBS Letters, 2006, 580 (13):3136–3144.

[21] Ko J H, Ham K H, Park S, et al, Plant body weight-induced secondary growth in Arabidopsis and its transcription phenotype revealed by whole-transcriptome profiling[J].Plant Physiol,2004,135(2): 1069–1083.

[22] Shin Y K, Yum H, Kim E S, et al. BcXTH1, a Brassica campestrishomologue of ArabidopsisXTH9,is associated with cell expansion[J]. Planta, 2006, 224(1):32–41.

[23] Hiwasa K, Nakano R, Inaba A, et al. Expression analysis of genes encoding xyloglucan endotransglycosylase during ripening in pear fruit[J]. Acta Horticulture,2003,628:549–553.

[24] Asif MH, Lakhwani D, Pathak S, et al. Transcriptome analysis of ripe and unripe fruit tissue of banana identifies major metabolic networks involved in fruit ripening process[J]. BMC Plant Biol, 2014,14: 316

[25] Cho SK, Kim JE, Park JA, et al. Constitutive expression of abiotic stress-inducible hot pepper CaXTH3, which encodes a xyloglucan endotransglucosylase/hydrolase homolog, improves drought and salt tolerance in transgenic arabidopsis plants. FEBS Lett, 2006, 580: 3136–3144

[26] Oogawara R, Satoh S, Yoshioka T, et al. Expression of alphaexpansin and xyloglucan endotransglucosylase /hydrolase gene associated with shoot elongation enhanced by anoxia, ethylene and carbon dioxide in arrowhead(Sagittaria pygmaea Miq.)tubers[J].Ann Bot,2005, 96:693–702.

[27] Eklof J M, Brumer H. The XTH gene family: an update on enzyme structure, function, and phylogeny in xyloglucan remodeling[J].Plant Physiol, 2010,153 (2): 456–466.

[28] Ookawara R, Satoh S, Yoshioka T, et al. Expression of α-expansin and xyloglucan endotransglucosylase/hydrolase genes associated with shoot elongation enhanced by anoxia, ethylene and carbon dioxide in arrowhead (Sagittaria pygmaea miq.) tubers. Ann Bot, 2005, 96: 93–702

[29] Osato Y, Yokoyama R, Nishitani K. A principal role for AtXTH18 in arabidopsis thaliana, root growth: a functional analysis using rnai plants. J Plant Res, 2006, 119: 153–162

[30] Genovesi V, Fornale S, Fry SC, et al. ZmXTH1, a new xyloglucan endotransglucosylase/hydrolase in maize, affects cell wall structure and composition in Arabidopsis thaliana[J]. Journal of Experimental Botany,2008, 59(4): 875–889.

[31] Han Y, Ban Q, Li H, et al. DkXTH8, a novel xyloglucan endotransglucosylase/hydrolase in persimmon, alters cell wall structure and promotes leaf senescence and fruit postharvest softening. Sci Rep, 2006, 6: 39155

[32] Harada T, Torii Y, Morita S, et al. Cloning, characterization,and expression of xyloglucan endotransglucosylase/hydrolase and expansin genes associated with petal growth and development during carnation flower opening. J Exp Bot, 2011, 62: 815–823

[33] Han Y, Han S, Ban Q, et al. Overexpression of persimmon dkxth1, enhanced tolerance to abiotic stress and delayed fruit softening in transgenic plants. Plant Cell Rep, 2017, 36: 583–596

[34] Siyan Liu; Jinfeng Liu; Yuzhe Zhang; Yushi Jiang; Shaowang Hu; Andi Shi; Qiyao Cong; Shuyan Guan; 2; Jing Qu; Yao Dan, Cloning of the Soybean sHSP26 Gene and Analysis of Its Drought Resistance,[J] Phyton (Buenos Aires) 2022.91(7):1465–1482

[35] Zhang Ye; Zhang Han zhu; Fu Jia yu; Du Ye yao; Qu Jing; Song Yang; Wang Pi wu, The GmXTH1 gene improves drought stress resistance of soybean seedlings,[J] Molecular Breeding, 2021, 42(1):14–21.

[36] Aranjuelo I, Molero G, Erice G et al. Plant physiology and proteomics reveals the leaf response to drought in alfalfa (Medicago sativa L.). J Exp Bot 2011, 62(1):111–123.

[37] Gall LH, Philippe F, Domon JM, et al. Cell wall metabolism in response to abiotic stress. Plants, 2015, 4: 112–166.

[38] Han WL, Kim J. Expansina 17 up-regulated by lbd18/asl20 promotes lateral root formation during the auxin response. Plant Cell Physiol, 2013, 54: 1600–1611.

[39] Baier, M.C., Barsch, A., Küster, H., and Hohnjec, N.. Antisense repression of the Medicago truncatula nodule-enhanced sucros synthase leads to a handicapped nitrogen fixation mirrored by specific alterations in the symbiotic transcriptome and metabolome. Plant Physiol. 2007, 145: 1600–1618.

[40] Olsen S, Striberny B, Hollmann J, et al. Getting ready for host invasion: elevated expression and action of xyloglucan endotransglucosylases/hydrolases in developing haustoria of the holoparasitic angiosperm cuscuta. J Exp Bot, 2015, 258: 193–204

[41] Blum, A. Osmotic adjustment is a prime drought stress adaptive engine in support of plant production [J]. Plant Cell and Environment, 2016,40(1): 4–10

[42] Chaves M M, Flexas J, Pinheiro C. Photosynthesis under drought and salt stress: regulation mechanisms from whole plant to cell[J]. Annals of Botany, 2008,103(4):551–560.

[43] Chen Q, Tao S Y, Bi X H, Xu X, Wang L L, Li SM. Research molecular biology mechanism of drought resistance in rice [J]. American Journal of Molecular BIOLOGY, 2015,03(2):102–107.

